# Development of autotrophy in *Escherichia coli* through adaptive laboratory evolution

**DOI:** 10.64898/2026.02.23.707615

**Authors:** Shao-Yu Huang, Jian-Hau Peng, Shou-Chen Lo, Chia-Ho Liu, Yu-Hsi Lin, En-Pei Isabel Chiang, Chieh-Chen Huang

**Affiliations:** Doctoral Program in Department of Life Sciences, National Chung Hsing University, Taichung City, 402, Taiwan; Doctoral Program in Microbial Genomics, National Chung Hsing University and Academia Sinica, Taichung City and Taipei City, 402 and 115, Taiwan; Department of Food Science and Biotechnology, National Chung Hsing University, Taichung City, 402, Taiwan; Advanced Plant and Food Crop Biotechnology Center, National Chung Hsing University, Taichung City, 402, Taiwan; Innovation and Development Center of Sustainable Agriculture, National Chung Hsing University, Taichung City, 402, Taiwan

## Abstract

Enabling heterotrophic *Escherichia coli* to use CO_2_ as its only carbon source remains a great challenge, and previous studies approached autotrophy conversion by metabolic engineering. Although its native carbon fixation routes were identified, the potential to reach autotrophy by itself has long been overlooked. In this study, autotrophy in *E. coli* was developed through adaptive laboratory evolution. After 1,000 days of consecutive inorganic subculturing, missense mutations were found in isocitrate dehydrogenase *icd* and isocitrate dehydrogenase kinase/phosphatase *aceK* genes, determining the metabolic switch between the citrate cycle and the glyoxylate shunt. By transcriptomic comparison of the adapted *E. coli* between inorganic and organic cultivations, two CO_2_ fixing enzymes activated in autotrophic mode were found, including the upregulated pyruvate:ferredoxin oxidoreductase YdbK and phosphoenolpyruvate carboxykinase Pck. Connected by the upregulated phosphoenolpyruvate synthase PpsA, a carbon fixation module was constituted, which was the shared foundation of the aspartate-threonine cycle and the citrate-glyoxylate-methylcitrate cycle, and thus integrating into an autotrophic network. By comparing the ^13^C enrichment patterns in inorganic cultivations between the adapted and initial *E. coli*, the favorable direction of the autotrophic network was confirmed.

**IMPORTANCE:** This is the first study to accomplish autotrophy in *E. coli* through long-term evolution alone. Besides missense mutations in *icd* and *aceK* genes, adapted *E. coli* also actively regulated its gene expression to respond to inorganic environment, such as directing the metabolic switch towards the glyoxylate shunt. For biomass formation, a carbon fixation module consisted of the upregulated YdbK, PpsA, and Pck produced pyruvate and oxaloacetate as precursors for two cycles. The aspartate-threonine cycle with a replenishment side loop further accumulated these precursors, and the citrate-glyoxylate-methylcitrate cycle was driven by four overexpressed enzymes to catalyze six reactions. These metabolic pathways were integrated into a novel autotrophic network, and by understanding the nature of *E. coli*, rational designs for its carbon fixation optimization become attainable by using compatible mechanisms.

## INTRODUCTION

To limit global warming to 1.5°C above pre-industrial levels with 50 % likelihood, the remaining carbon budget in 2025 was estimated to be 130 gigatons of CO_2_ (GtCO_2_), which will soon be exhausted if the emission rates stay at 2024 levels, namely 42 GtCO_2_ (1). As a sustainable way of carbon reduction, metabolic engineering can replace high carbon-emitting manufacturing. Although there are autotrophs that can already fix CO_2_ efficiently, their applications are hindered by the difficulties in cultivation, bioproduction, and genetic engineering (2). In contrast, *Escherichia coli* is a well-studied model organism that is categorized as a strict heterotroph, which is widely used due to its rapid growth in diverse conditions, industrial scalability, and the ease of genetic engineering (2, 3). Therefore, its trophic conversion to autotrophy remains a valuable yet challenging goal.

In a previous study of constructing synthetic autotrophy in *E. coli* (4), the Calvin cycle was chosen due to its ubiquity in the biosphere (5). Before introducing the Calvin cycle, the native pathways that would disrupt its operation were blocked by deleting three genes, including phosphofructokinases (*pfkA/B*) and glucose-6-phosphate dehydrogenase (*zwf*). To introduce the Calvin cycle and its supporting mechanisms, four heterologous genes were used, including ribulose-1,5-bisphosphate carboxylase/oxygenase RuBisCO (*cbbM*) to fix CO_2_, phosphoribulokinase (*prkA*) to connect the Calvin cycle, carbonic anhydrase CA (*Rru_A2056*) to increase CO_2_ concentration, and formate dehydrogenase (*fdh*) to generate NADH by oxidizing formate supplement. Lastly, to allow beneficial mutations and metabolic rewiring to happen, adaptive laboratory evolution (ALE) was conducted, along with xylose supplement to favor the operation of the Calvin cycle. After 343 days of ALE, xylose supplement was deprived, and the evolved autotrophic strains were selected. In their accumulated mutations, some are related to the branch points of the Calvin cycle, and the reduced or abolished activities of their encoded enzymes would help stabilizing its operation. In a follow-up study (6), three mutations were sufficient to enable autotrophy in the engineered *E. coli*.

Another study focused on using the native carbon fixation route in *E. coli* to approach synthetic autotrophy (7). By computational analysis, three routes were identified, including the Gnd-Entner-Doudoroff (GED) cycle, the reductive glycine (rGly) pathway, and the serine-threonine (Ser-Thr) cycle. Among them, the GED cycle was chosen due to its fewer required reactions and more oxygen tolerance. To block the undesired pathways, six disruptive genes were deleted, including *pfkA/B*, *zwf*, fructose-6-phosphate aldolases (*fsaA/B*), and 1-phosphofructokinase (*fruK*). To enforce the GED cycle, three endogenous genes were overexpressed to compensate their natively low expression levels, including 6-phosphogluconate dehydrogenase (*Gnd*) to enhance CO_2_ fixation, and phosphogluconate dehydratase (*Edd*) and 2-keto-3-deoxygluconate 6-phosphate aldolase (*Eda*) to sustain the GED cycle. After 6–8 days of short-term evolution with xylose supplement, the growth facilitated by the GED cycle was observed. Despite its growth dependence on xylose supplement, if longer evolution time was given, it might be gradually reduced and thus approaching fully autotrophic growth.

Despite there are precedents of synthetic autotrophy in *E. coli*, their ways of living were predetermined by experimental designs. Decisions on the target pathways were based on their theoretical feasibilities and efficiencies, which could be incompatible with the native metabolism of *E. coli*. Deletions of disruptive genes shaped the native metabolism for the target pathways to fit in, and gene introduction or overexpression enforced the target pathways, yet these genetic manipulations could have profound and unknown consequences. While ALEs allowed *E. coli* to explore its potential, the organic supplements provided selective advantages to the target pathways, which in turn narrowed the diversity of evolution, and unforeseen pathways could be lost along the process. All these designs were derived from the assumption that *E. coli* cannot reach autotrophy by itself. However, the discovery of the native carbon fixation routes in *E. coli* (7) raised the possibilities for its autotrophy to develop, which should be considered and yet have long been overlooked. Through evolution, unexpected abilities might emerge for *E. coli* to gain fitness (8, 9). Therefore, our study aimed to develop autotrophy in *E. coli* by ALE alone, reducing human interferences for its natural tendency to unfold. After gaining systematic views on how *E. coli* reaches autotrophy, optimization of its carbon fixation could be rationally designed in the future.

## RESULTS

### Growth in different ALE phases

At the first attempt, the initial *E. coli* BW25113 with plasmid pGETS118 providing antibiotic resistance was cultivated in inorganic M9 medium. After three days, its population dropped below 3% of the inoculated number. To avoid extinction before *E. coli* developed the ability to grow autotrophically, the first phase of ALE was carried out by inorganic-organic rotation. For one rotation, the remaining culture after three days of inorganic cultivation was transferred to organic M9 medium with glucose, then the repopulated culture was reintroduced to inorganic M9 medium, cultivating for another three days (Fig. 1A). After 27 cumulative days of inorganic cultivation, the preliminarily adapted *E. coli* was able to double the inoculated number, which was the first sign of growth. After 153 cumulative days of inorganic cultivation, the growth of the preliminarily adapted *E. coli* increased to above 10 folds of the inoculated number (Fig. 1B).

**Fig 1.**
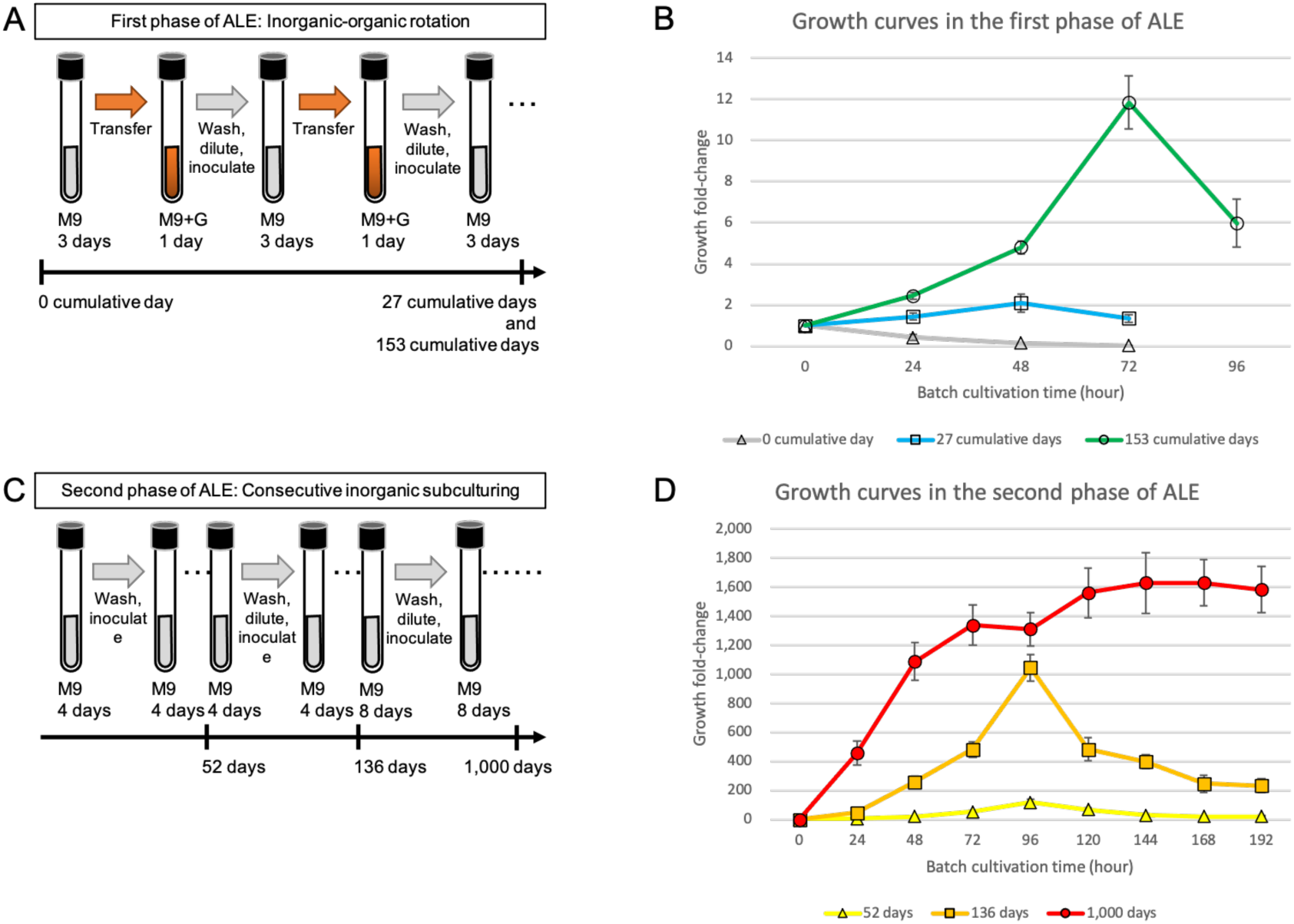
ALE experiments in different phases, and their growth curves. (A) Workflow of ALE in the first phase, and the cumulative days of inorganic cultivation for *E. coli* batches (black arrow). (B) Growth curves in the first phase of ALE, including *E. coli* batches that were inorganically cultivated for 0, 27, and 153 cumulative days. (C) Workflow of ALE in the second phase, and the days of consecutive inorganic subculturing for *E. coli* batches (black arrow). (D) Growth curves in the second phase of ALE, including *E. coli* batches that were subcultured for 52, 136, and 1,000 days in this phase. Data shown are the growth curves with the inoculated numbers between 1,500–2,500 CFU/mL. Growth fold-changes are calculated by dividing CFU/mL at hour N by the inoculated CFU/mL of that batch, and their means ± standard deviations of six biological replicates are depicted. Acronyms: M9, inorganic M9 medium; M9+G, organic M9 medium with 0.1% glucose.

To ensure such growth was not enabled by utilizing nutrients that were stored during organic cultivation, the second phase of ALE was carried out by consecutive inorganic subculturing, avoiding the involvement of organic M9 medium (Fig. 1C). After 52 days, the autotrophic growth of the adapted *E. coli* improved to above 100 folds. After 136 days, its autotrophic growth surpassed 1,000 folds. After 1,000 days, its autotrophic growth exceeded 1,600 folds, and the declination of its population was alleviated, showing improved longevity in stationary phase (Fig. 1D).

### Mutations and their relations with autotrophy

Autotrophic growth could be enabled by mutations accumulated during ALE. To examine their relations, five adapted clones after 1,000 days of consecutive inorganic subculturing were sequenced, and 101 mutations were identified (Table 1; Table S1).

**Table 1.** Mutation types and numbers.

| Mutation type | Number |
| --- | --- |
| Plasmid non-coding region mutation | 29 |
| Genome non-coding region mutation | 21 |
| Genome stop-codon-loss mutation | 1 |
| Genome stop-codon-gain mutation | 3 |
| Genome synonymous mutation | 20 |
| Genome missense mutation | 27 |
| Total | 101 |

There were 29 mutations in plasmids, all occurred in non-coding regions, distancing over 400 bp from their downstream genes (Table S1). As the plasmid only provided antibiotic resistance to prevent contamination during cultivation, and no autotrophic mechanism was added, these mutations are unrelated to autotrophy. There were also 21 non-coding region mutations in genomes, but their closest downstream genes are unrelated to the central carbon metabolism (Table S1), making the roles of these non-coding region mutations unclear.

There were four mutations that changed protein length, including one stop-codon-loss mutation and three stop-codon-gain mutations. The only stop-codon-loss mutation could be found in all clones, causing the extension of a fimbrial-like protein SfmA (*181Glnext*42) (Table S1). However, it is >99% identical with a fimbrial subunit FimA in *E. coli*, hence this stop-codon-loss mutation might be unrelated to autotrophy. In contrast, all three stop-codon-gain mutations would cut off the syntheses of functional regions, causing loss-of-functions of the affected proteins, including aerobic respiration two-component sensor histidine kinase ArcB (Glu510*) (10), ethanol dehydrogenase EutG (Cys91*), and flagellar hook-filament junction FlgL (Gly156*) (Table S1). However, all stop-codon-gain mutations occurred in the same clone, and such infrequency suggests that these stop-codon-gain mutations were nonessential for autotrophy.

There were 27 missense mutations, and only five of them occurred in all clones, altering amino acids in ArcB (Asp166Gly), fatty acid metabolism transcriptional regulator FadR (Asp90Glu), glutamine ABC transporter substrate-binding protein GlnH (Ala17Val), exoribonuclease II Rnb (Val278Leu), and uncharacterized lipoprotein YdcL (Ile105Thr) (Table S1). However, none of these alterations located on known active sites. To focus on mutations that might be more relevant to autotrophy, two missense mutations related to the citrate cycle were identified. One of them affected isocitrate dehydrogenase Icd (Asp398Glu), which could be found in two clones (Table S1). Icd is responsible for the interconversion of isocitrate and *α*-ketoglutarate (AKG), and the altered amino acid Asp398 was close to NADP^+^ binding residues Arg395 and Tyr391(11). Additionally, within 20 synonymous mutations, nine of them occurred in *icd* gene (Table S1), indicating it could be a mutational hotspot during inorganic cultivation. Another citrate cycle related missense mutation affected Icd kinase/phosphatase AceK (Tyr474Ser), which could be found in four clones (Table S1). AceK can either deactivate Icd by phosphorylation, or activate Icd by dephosphorylation. The altered amino acid Tyr474 was next to Mg^2+^ binding residue Asp475, and Mg^2+^ is an essential cofactor for AceK to be functional (12, 13). Both Icd and AceK determine the metabolic switch between the citrate cycle and the glyoxylate shunt. During starvation, *E. coli* tends to bypass the citrate cycle through the glyoxylate shunt as a coping mechanism (14). Therefore, these two missense mutations that affected Icd (Asp398Glu) and AceK (Tyr474Ser) are most relevant to autotrophy.

Another aspect of mutation is the alteration of DNA methylation site. As an epigenetic mechanism, DNA methylation is a heritable and reversible modification that affects gene expression, especially when it occurs in a promoter region (15). There are two methyltransferases that have widespread sites in *E. coli* genome, including DNA adenine methyltransferase (Dam) that recognizes GATC consensus sequence, and DNA cytosine methyltransferase (Dcm) that recognizes CCAGG or CCTGG consensus sequences (16). While there was no mutation that affected Dcm sites, there were six mutations that either disrupting or creating Dam methylation site, including five site-disruption mutations and one site-creation mutation (Table S1). Among them, two site-disruption mutations were located in non-coding region, and both disrupted the same methylation site (17), distancing over 280 bp from a prophage side tail fiber encoding gene *stfR*, which is unrelated to the central carbon metabolism. Other four methylation site related mutations were located in coding regions, distancing over 270 bp from their promoter regions, and all of the potentially affected genes have no obvious relation with autotrophy.

### Differential expression of the native CO_2_ fixing routes

In addition to waiting for beneficial mutations to happen, the adapted *E. coli* could actively regulate its gene expression to respond to inorganic environment. By transcriptomic analysis of the adapted *E. coli* in inorganic and organic cultivations, the expression levels of the carbon fixation enzymes were compared (Table 2). By identifying the upregulated carbon fixing enzymes, their downstream metabolisms could be the pathways taken by the adapted *E. coli* to reach autotrophy. According to previous study, there are three native carbon fixation routes in *E. coli*, including the GED cycle, the rGly pathway and the Ser-Thr cycle (7). The GED cycle starts with the reductive carboxylation of 6-phosphogluconate (6PG) to ribulose 5-phosphate (Ru5P) by Gnd, which is the CO_2_ fixing enzyme of this cycle. Followed by two enzymes of the Entner-Doudoroff (ED) pathway, namely Edd and Eda, 6PG is converted to glyceraldehyde 3-phosphate (GAP) and pyruvate. Afterwards, pyruvate is processed through gluconeogenesis to produce GAP, and the accumulated GAPs are processed through the pentose phosphate pathway for Ru5P replenishment, and thus starting a new round of the GED cycle. In our results, the expression levels of Gnd, Edd, and Eda were downregulated in autotrophic mode (Table 2; Table S2), showing that the GED cycle was not activated by the adapted *E. coli* when facing inorganic adversity. In previous study, other than compensating the natively low expression levels of Gnd, Edd, and Eda by overexpressing them, the growth of their engineered *E. coli* was dependent on xylose supplement (7), which served as the precursor of Ru5P, indicating that the replenishment of Ru5P was an unsolved problem for the GED cycle to enable fully autotrophic growth.

**Table 2.** Differential expression of the carbon fixing enzymes. Expression fold-changes of the adapted E. coli in inorganic cultivation relative to in organic cultivation are shown.

| EC number | Enzyme name | Enzyme description | Expression fold-change |
| --- | --- | --- | --- |
| 1.1.1.44 | Gnd | 6-phosphogluconate dehydrogenase | 0.073 |
| 1.4.4.2 | GcvP | Glycine decarboxylase | 0.088 |
| 1.2.7.1 | YdbK | Pyruvate:ferredoxin oxidoreductase | 1.786 |
| 4.1.1.49 | Pck | Phosphoenolpyruvate carboxykinase | 2.358 |
| 4.1.1.31 | Ppc | Phosphoenolpyruvate carboxylase | 0.368 |
| 1.1.1.38 | MaeA | NAD-dependent malic enzyme | 0.506 |

The rGly pathway begins with the tetrahydrofolate-mediated one-carbon (THF-C1) metabolism of formate, and the accumulated 5,10-methylene-THFs participate in two reactions. One 5,10-methylene-THF is condensed with CO_2_ and ammonia to produce glycine, which is carried out by the enzymes of the glycine cleavage system (GCS), including glycine decarboxylase (GcvP) as the CO_2_ fixing enzyme of the rGly pathway (18). Another 5,10-methylene-THF is condensed with the produced glycine to serine by serine hydroxymethyltransferase (GlyA), and the produced serine is then converted to pyruvate by five serine deaminases (19). In our results, the GCS enzymes were downregulated (Table 2; Table S3). The problem of the rGly pathway could be the replenishment of 10-formyl-THF, which is the precursor of 5,10-methylene-THF. As the hydrolysis reaction by 10-formyl-THF deformylase (PurU) is irreversible (20), and phosphoribosylglycinamide formyltransferases (PurN/T) that produce 10-formyl-THF by utilizing formate were downregulated, the replenishment of 10-formyl-THF was unsustainable, leading to an infeasible THF-C1 metabolism, and thus disconnecting the rGly pathway.

In the Ser-Thr cycle, glycine still needs to be converted to serine by GlyA, involving the infeasible THF-C1 metabolism to provide 5,10-methylene-THF. After the deamination of serine, the produced pyruvate is phosphorylated to phosphoenolpyruvate (PEP) by PEP synthetase (PpsA). Then, PEP is carboxylated to oxaloacetate (OAA) by PEP carboxykinase (Pck), which is the CO_2_ fixing enzyme of this cycle (21, 22). After the processes of aspartate biosynthesis, threonine biosynthesis, and threonine degradation, glycine and acetyl-CoA are produced. To solve the problem of the infeasible THF-C1 metabolism, pyruvate:ferredoxin oxidoreductase (YdbK) (23, 24) could directly bridge acetyl-CoA to pyruvate by reductive carboxylation, simultaneously fixing CO_2_ and bypassing the infeasible THF-C1 metabolism. By connecting YdbK, PpsA, and Pck, a carbon fixation module is formed, which is the foundation of the aspartate-threonine (Asp-Thr) cycle (Fig. 2). While acetyl-CoA connects the Asp-Thr cycle, glycine can either directly serve as a building block for protein synthesis, or be repurposed through deamination to glyoxylate by D-amino acid dehydrogenase (DadA) (25). After two molecules of glyoxylate are accumulated by fixing four CO_2_, the condensation by glyoxylate carboligase (Gcl) produces tartronate semialdehyde, along with the leakage of one CO_2_. As the amount of fixed CO_2_ exceeds the leaked CO_2_, it remains a feasible autotrophic pathway. Tartronate semialdehyde is then converted to glycerate by tartronate semialdehyde reductase (GarR), and glycerate is converted to 2-phosphoglycerate (2PG) by glycerate 2-kinase (GarK), and thus connecting the glycerate pathway (26). Finally, 2PG can be further converted to PEP by enolase (Eno), resulting in a replenishment side loop of the Asp-Thr cycle (Fig. 2). In our results, YdbK, PpsA, and Pck as the enzymes of the carbon fixation module were upregulated (Table 2; Table S4). In contrast to the downregulated CO_2_ fixing enzymes of the GED cycle and the rGly pathway, the Asp-Thr cycle could be more feasible due to its activated carbon fixation module.

**Fig 2.**
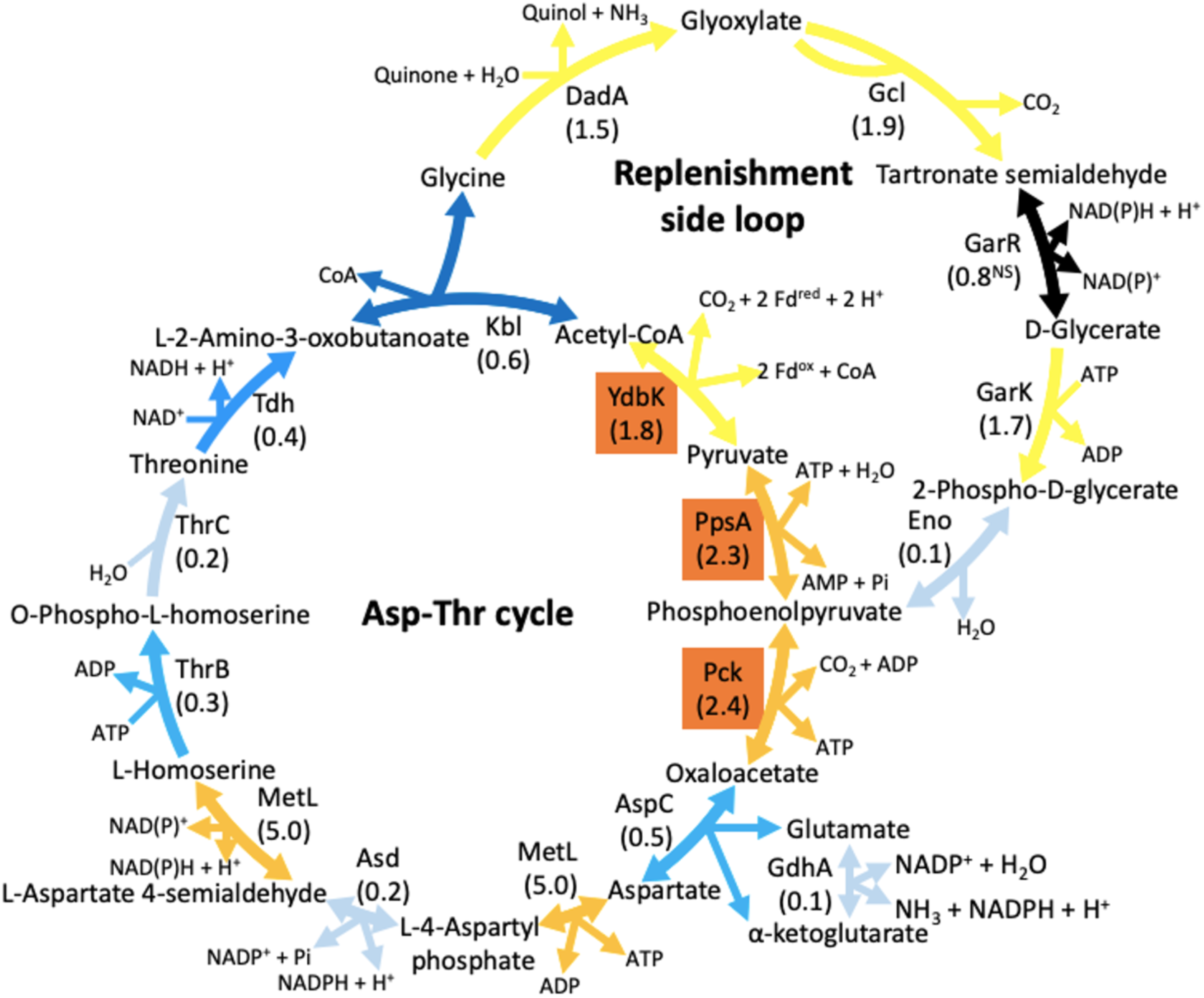
Differential expression of the Asp-Thr cycle in autotrophic mode. The Asp-Thr cycle consists of the carbon fixation module with three enzymes (orange boxes), aspartate synthesis, threonine synthesis, threonine degradation, and a glycerate pathway as the replenishment side loop. Numbers in parentheses represent expression fold-changes of the adapted *E. coli* in inorganic cultivation relative to organic cultivation. NS symbol represents the enzyme with expression fold-change that is statistically non-significant (q-value > 0.05) of two biological replicates. Acronyms: Fd^red^, reductive ferredoxin; Fd^ox^, oxidative ferredoxin.

### Differential expression of the citrate cycle related metabolisms

Pyruvate and OAA produced by the carbon fixation module are not only the components of the Asp-Thr cycle, but also the upstream metabolites of the citrate cycle, which is responsible for the syntheses of other core metabolites, such as AKG and succinyl-CoA (27). In the citrate cycle, there are other carbon fixing enzymes, including PEP carboxylase (Ppc) and two malate dehydrogenases (MaeA/B) (21). In our results, however, Ppc and MaeA were downregulated, and MaeB was slightly upregulated without statistical significance (Table 2; Table S5). Therefore, the upregulated YdbK and Pck were the main CO_2_ fixing enzymes for the adapted *E. coli* in autotrophic mode.

When entering the citrate cycle, OAA could either be reduced to malate by malate dehydrogenase (Mdh), or be condensed with acetyl-CoA to citrate by citrate synthase (GltA) and the bifunctional methylcitrate synthase (PrpC) (28). In our results, Mdh and GltA were downregulated, whereas PrpC was overexpressed (Table S5), suggesting that OAA tended to be converted to citrate by PrpC. Following such tendency, citrate is then isomerized to isocitrate, catalyzing by aconitate hydratase B (AcnB) and the bifunctional methylcitrate dehydratase (PrpD) (29). In our results, AcnB was upregulated, and PrpD was overexpressed (Fig. 3; Table S5), supporting the metabolic tendency towards PrpC and PrpD. Isocitrate is the branch point metabolite between the citrate cycle and the glyoxylate shunt, and the metabolic switch is determined by Icd and AceK. If isocitrate was directed towards the citrate cycle, it would be decarboxylated to AKG by Icd, along with the leakage of CO_2_. If isocitrate was directed towards the glyoxylate shunt, Icd would be deactivated by the upregulated AceK (30, 31), and isocitrate would be split into glyoxylate and succinate by isocitrate lyase (AceA). In our results, Icd was downregulated, whereas AceK and AceA were overexpressed (Table S5), suggesting that the metabolic switch was directed towards the glyoxylate shunt, and thus conserving the fixed carbon by reducing the leaked CO_2_. Nevertheless, the decarboxylation by Icd cannot be bypassed completely due to the necessity to produce AKG, which is the essential precursor for glutamate and other derived amino acids. As a common way to cope with starvation (14), the glyoxylate shunt is often coupled with the recycling of acetate (32), catalyzing by acetyl-CoA synthetase (Acs) and the bifunctional propionate CoA ligase (PrpE) (33). In our results, Acs and PrpE were overexpressed (Table S5), suggesting that the acetate recycling was also activated in autotrophic mode.

**Fig 3.**
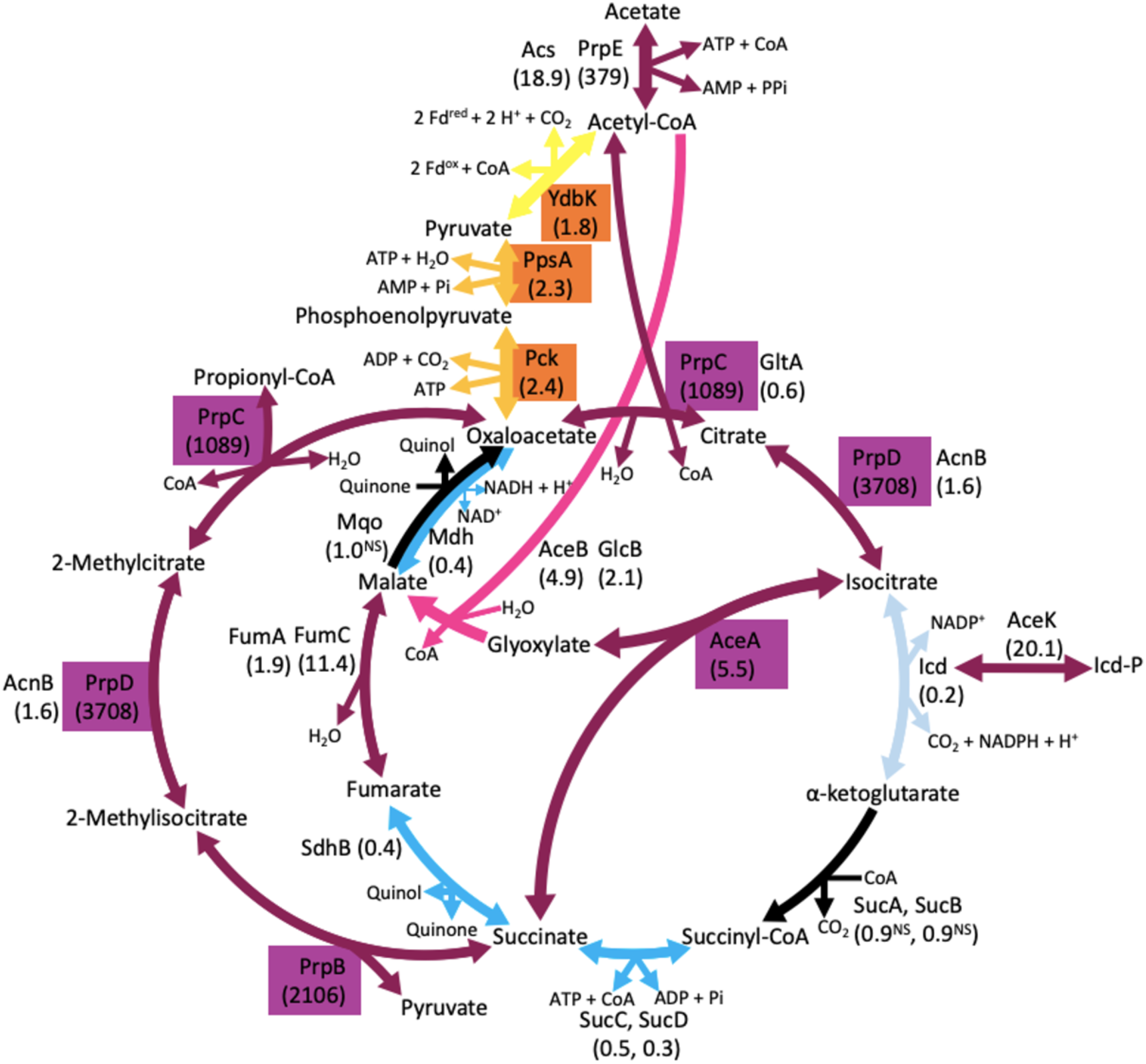
Differential expression of the citrate cycle related metabolisms in autotrophic mode. The enzymes that constitute the carbon fixation module (orange boxes) and the CGM cycle (purple boxes) are shown. Numbers in parentheses represent expression fold-changes of the adapted *E. coli* in inorganic cultivation relative to organic cultivation. NS symbols represent the enzymes with expression fold-changes that are statistically non-significant (q-value > 0.05) of two biological replicates. Acronyms: Icd-P, phosphorylated Icd (deactivated); Fd^red^, reductive ferredoxin; Fd^ox^, oxidative ferredoxin.

As the products of isocitrate splitting, glyoxylate and succinate can be further processed in several ways. For glyoxylate, it either joined in the replenishment side loop of the Asp-Thr cycle by the upregulated Gcl (Fig. 2), or merged back to the citrate cycle by condensing with acetyl-CoA to produce malate, catalyzing by malate synthase A (AceB) and malate synthase G (GlcB). In our results, GlcB was upregulated, and AceB was overexpressed (Table S5). Afterwards, malate can either be oxidated to OAA by Mdh and malate:quinone oxidoreductase (Mqo) (34), or be dehydrated to fumarate by five independent fumarases (FumA/B/C/D/E). In our results, Mdh was downregulated, Mqo was slightly downregulated without statistical significance, whereas FumA were upregulated, and FumC was overexpressed (Table S5), suggesting that malate tended to be converted to fumarate. Another product of isocitrate splitting is succinate, which could merge back to the citrate cycle, either by condensing with CoA to produce succinyl-CoA by succinyl-CoA synthetase complex (SucCD), or oxidizing to fumarate by succinate dehydrogenase complex (SdhABCD) and fumarate reductase complex (FrdABCD). In our results, however, SucCD, SdhB, and FrdABCD were downregulated (Table S5), and the downregulation of any subunit would lead to the decreased amount of functional complete complex, suggesting that these functions were not activated in autotrophic mode. An alternative way to process succinate is through the reversed methylcitrate pathway (35), starting by condensing with pyruvate to methylisocitrate by methylisocitrate lyase (PrpB). Afterwards, methylisocitrate was isomerized by the bifunctional PrpD to produce methylcitrate, which was then split into propionyl-CoA and OAA by the bifunctional PrpC. Propionyl-CoA is the precursor of isoleucine (36) and fatty acids (37), and OAA can merge to the Asp-Thr cycle (Fig. 2) or the citrate cycle. In our results, PrpBCD were overexpressed (Table S5), suggesting the feasibility of the reversed methylcitrate pathway. By connecting six reactions that are carried out by only four overexpressed enzymes, namely PrpBCD and AceA, a citrate-glyoxylate-methylcitrate (CGM) cycle is formed (Fig. 3). As the CGM cycle and the Asp-Thr cycle are rooted from the same carbon fixation module, these metabolic pathways can be integrated into an autotrophic network (Fig. 4).

**Fig 4.**
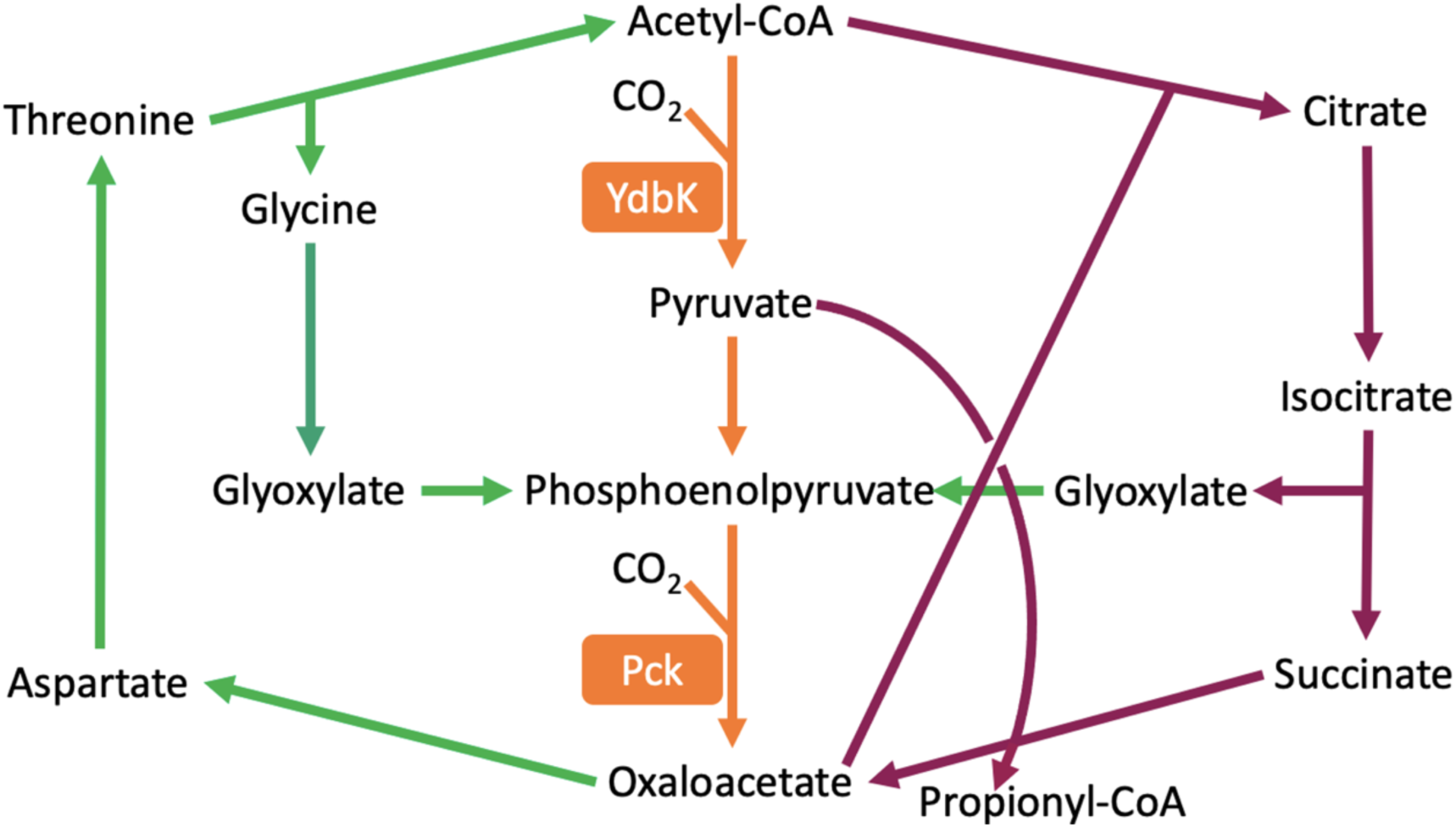
Autotrophic network. The integration of the carbon fixation module (orange arrows and boxes), the Asp-Thr cycle (green arrows), and the CGM cycle (purple arrows).

### Differential expression of the iron-sulfur cluster synthesis

In the autotrophic network, many reactions are carried out by metalloproteins, which require iron-sulfur (FeS) clusters to be functional (38), including CO_2_ fixation by YdbK, electron transfer by monomeric ferredoxins, and aconitate synthesis by AcnB and PrpD. Therefore, FeS cluster plays an essential role in autotrophy.

Before synthesizing FeS clusters, iron and sulfur ions need to be transported into cell bodies. For iron uptake, other than passive diffusion through outer membrane porins (OmpC/F), there are six active transport systems, including ferrous (Feo), ferric dicitrate (Fec), enterobactin (Ent), ferrichrome (Fhu), ferric iron uptake (Fiu), and colicin I receptor (Cir) systems (39–41). In our results, except for OmpC and Feo system, most proteins from other five systems were upregulated in autotrophic mode (Table S6). For iron storage, ferritin related proteins (FtnAB) were downregulated, and bacterioferritin (Bfr) had non-significant change (Table S6). Due to the abundance of Fe^2+^ supplement in inorganic M9 medium, iron storage as a response to iron deficiency was unnecessary. For sulfur uptake, there are sulfate/thiosulfate (CysPUWA/Sbp), alkanesulfonate (SsuACB), and taurine (TauACB) transport systems. Followed by serial reduction and cysteine synthesis, the sulfur metabolism is completed, and the produced sulfide and cysteine can be used for FeS cluster assembly (42). In our results, except for TauACB, most proteins were upregulated (Table S7).

For FeS cluster assembly, there are two independent systems (43), namely the iron-sulfur cluster (Isc) system that is encoded by the *iscRSUA-hscBA-fdx-iscX* operon (44) and the sulfur mobilization (Suf) system that is encoded by the *sufABCDSE* operon (45, 46). In our results, proteins of the Suf system were either upregulated or overexpressed, whereas IscR as the *isc* operon repressor was upregulated (47), and most proteins from the Isc system were downregulated (Table S8). Due to the oxygen tolerance of the Suf system (48, 49), it could be the main assembly mechanism under aerobic conditions. Afterwards, the assembled FeS clusters are delivered to target apoproteins, resulting in 144 annotated metalloproteins that are responsible for electron transfer, radical generation, gene regulation, and metabolite synthesis (38).

As a form of reducing power, reductive ferredoxins drive the reductive carboxylation by YdbK, resulting in CO_2_ fixation. Among all monomeric ferredoxins (38), YgcO, and YfhL were upregulated, and Bfd was overexpressed (Table S9). Among the activated monomeric ferredoxins, YfhL has the reduction potential of -675 mV (50), which is strong enough to drive reductive carboxylation. Furthermore, the energy of reductive ferredoxins can trickle down to other forms of energy, starting with the conversion of reductive ferredoxin to NADPH by ferredoxin NADP^+^ reductase (Fpr). Afterwards, NADPH can be converted to NADH by soluble pyridine nucleotide transhydrogenase (SthA), and NADH can be converted to ATP through oxidative phosphorylation, and thus fulfilling the energy requirements of biomass production. In our results, however, only SthA and SdhC were upregulated (Table S9), and the latter is a subunit of the SdhABCD complex, along with the downregulated SdhB that limited the amount of functional complete complex. Such overall downregulation suggests that the redox balance in autotrophic mode is strictly controlled.

### ^13^C enrichment patterns of the citrate cycle metabolites

Although the autotrophic network was discovered by the activated enzymes, most of their responsible reactions are reversible, including two CO_2_ fixing enzymes. Such uncertainty could lead to carbon fixation or carbon loss as two opposite outcomes, challenging the foundation of autotrophy. Since the favorable directions of enzymes cannot be verified by differential expression alone, isotope labeling was needed to track the carbon flows in the adapted *E. coli*. Therefore, the ^13^CO_2_ supplemented inorganic cultivations of the adapted and the initial *E. coli* were conducted, and their ^13^C labeled metabolites from the citrate cycle were analyzed by GC-MS. By comparing their fractional ^13^C enrichment patterns, the favorable direction of the autotrophic network in the adapted *E. coli* could be clarified. Moreover, the ^13^C accumulated metabolites produced by the Asp-Thr cycle also participate in the citrate cycle (Fig. 5), hence the connections between these cycles could be observed.

**Fig 5.**
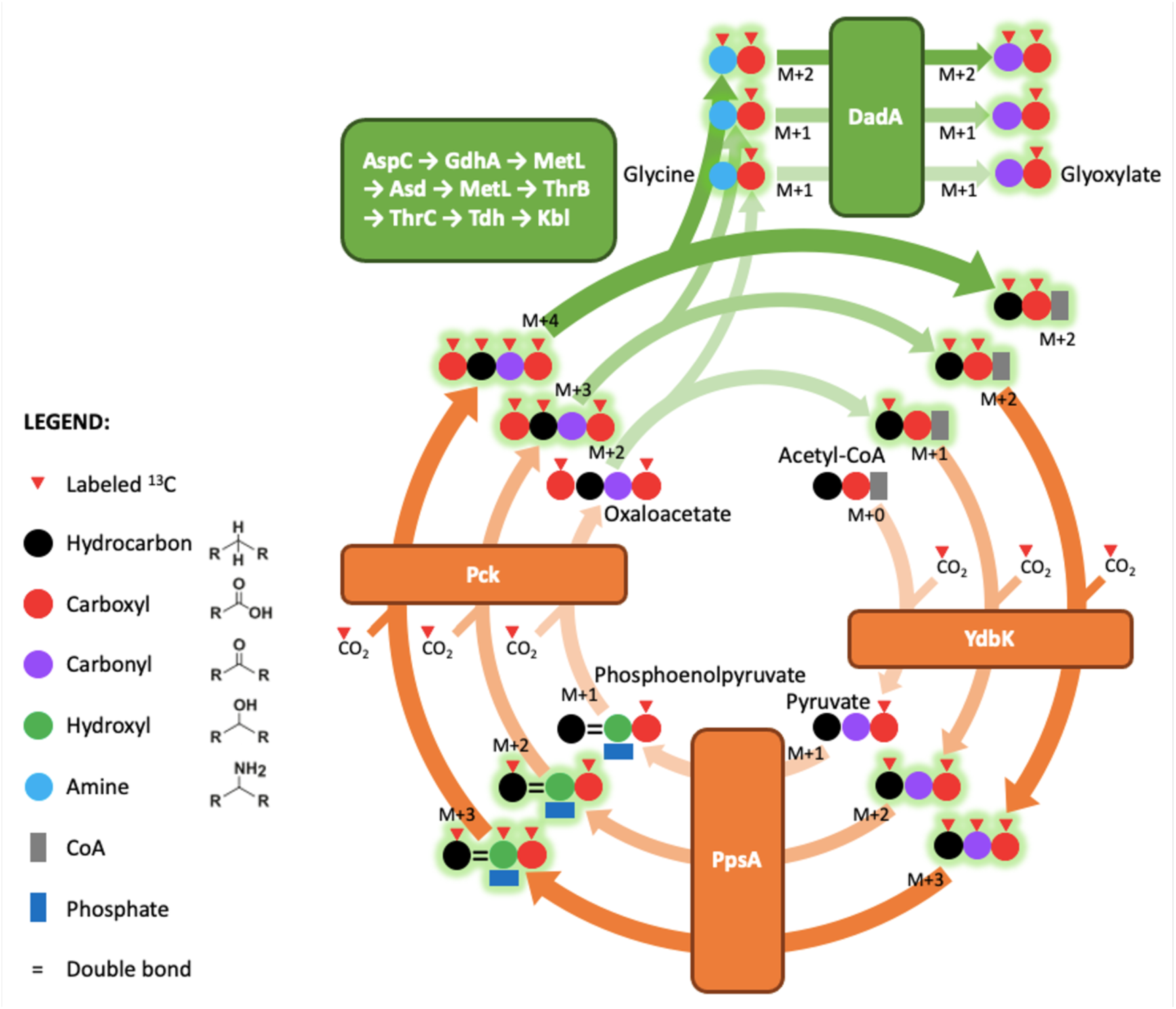
Asp-Thr cycle and its production of the ^13^C accumulated metabolites. The ^13^C atoms (red triangles) are incorporated by the carbon fixation module (orange arrows and boxes), and other processes (green arrows and boxes) are shown. Symbols and colors of metabolite carbons are illustrated in legend. The ^13^C accumulated metabolites (green shaded symbols) are produced by the Asp-Thr cycle, which can participate in the citrate cycle, and thus change the ^13^C enrichment patterns of the citrate cycle metabolites. Isotopolouges with different ^13^C enriched masses are denoted as M+*n* species.

As the first potential CO_2_ fixing enzyme of the carbon fixation module, YdbK carries out an oxidation-reduction reaction. If its favorable direction was towards reductive carboxylation, then ^13^CO_2_ would be fixed with acetyl-CoA, resulting in a mass (M) +1 pyruvate, which is an isotopologue labeled with one ^13^C atom. In our results, M+1 pyruvate was increased in the adapted *E. coli* (Fig. 6). Additionally, M+1 lactate was increased in the adapted *E. coli* (Fig. 6), which could be derived from the fermentation of the M+1 pyruvate (51). Both results confirm that YdbK favors reductive carboxylation, and thus enabling the first step of the carbon fixation module in the adapted *E. coli*.

**Fig 6.**
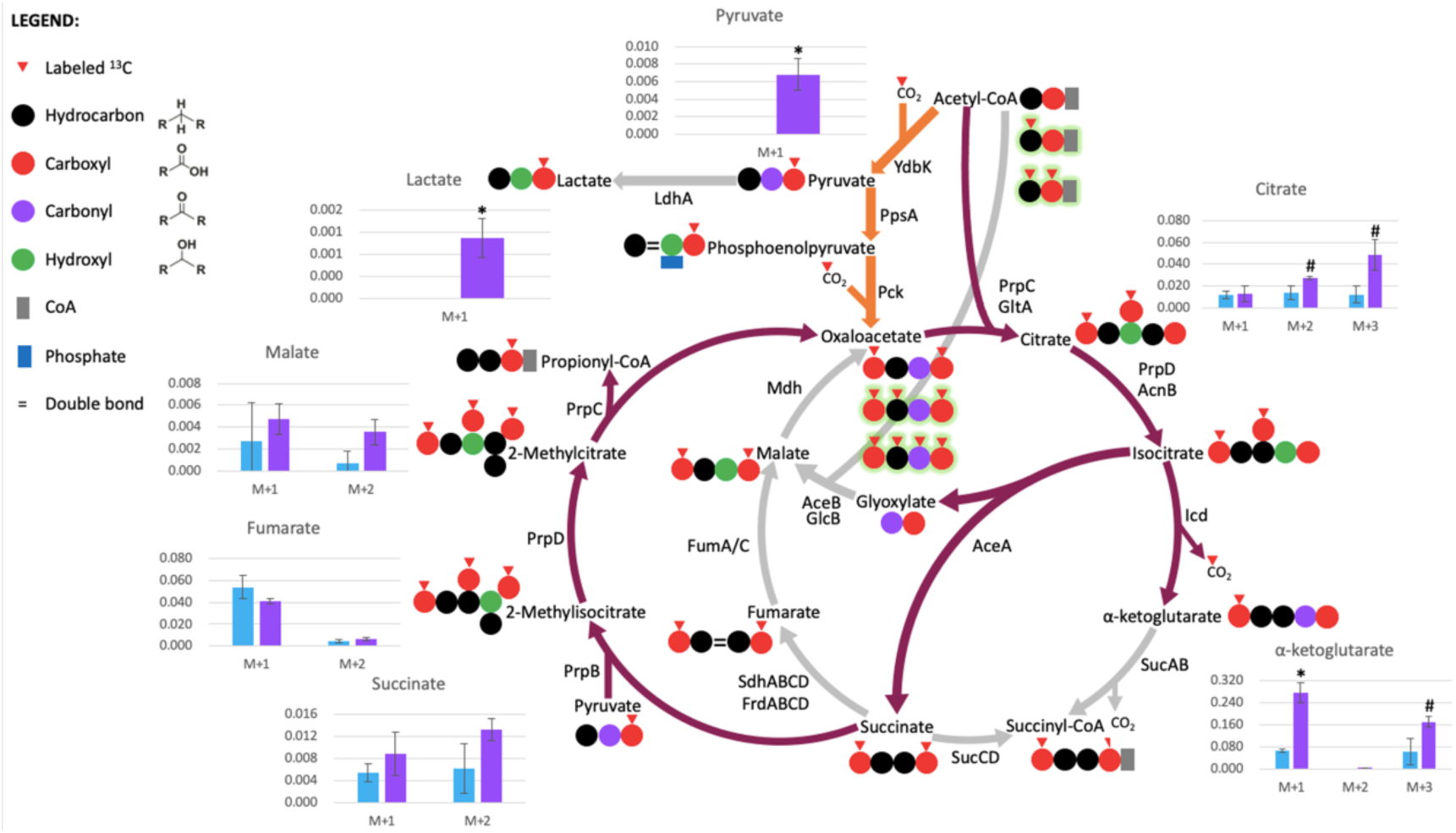
^13^C enrichment patterns of the citrate cycle related metabolites. The favorable directions of the CGM cycle reactions (purple arrows) are shown. Symbols and colors of metabolite carbons are illustrated in legend. ^13^C atoms (red triangles) and the ^13^C accumulated metabolites produced by the Asp-Thr cycle (green shaded symbols) would affect the ^13^C enrichment patterns of the citrate cycle metabolites. Bar graphs represent the fractional ^13^C enrichment patterns of metabolites, and the means ± standard deviations of the initial *E. coli* (light blue bars) and the adapted *E. coli* (light purple bars) are depicted. Isotopolouges with different ^13^C enriched masses are denoted as M+*n* species. Asterisks represent the means that are statistically significant (p-value < 0.05) of two biological replicates. Hashes represent the means that are borderline significant (p-value < 0.1) of two biological replicates.

Afterwards, M+1 pyruvate could be processed through PpsA and the second potential CO_2_ fixing enzyme Pck, resulting in the production of M+2 OAA. Due to the instability of OAA that makes it hard to detect (52), the production of M+2 OAA had to be inferred by the ^13^C enrichment patterns of its downstream metabolites. There are two ways to process M+2 OAA, namely the Asp-Thr cycle and the citrate cycle. When entering the Asp-Thr cycle, M+2 OAA leads to the production of the ^13^C accumulated metabolites (Fig. 5).

When entering the citrate cycle, M+2 OAA could either be converted to malate by Mdh, or be condensed with acetyl-CoA to citrate by PrpC. In our results, malate in M+1 and M+2 forms and citrate in M+1 form had non-significant changes, whereas citrate in M+2 and M+3 forms were increased in the adapted *E. coli* (Fig. 6). The ^13^C enrichment differences between malate and citrate show that M+2 OAA tended to be converted to citrate, which is consistent with PrpC overexpression and Mdh downregulation (Table S5). The increased M+2 citrate required the availability of its precursor M+2 OAA, inferring that Pck favored reductive carboxylation, and thus confirming the second step of the carbon fixation module in the adapted *E. coli*. The increased M+3 citrate required the participation of ^13^C accumulated metabolites, which were produced by the Asp-Thr cycle (Fig. 5), and thus implying the connection between the Asp-Thr cycle and the citrate cycle.

After citrate is isomerized by PrpD, the produced isocitrate could be processed either by the citrate cycle or the glyoxylate shunt. When following the citrate cycle, isocitrate is decarboxylated to AKG by Icd, causing the leakage of one CO_2_. According to the downregulated Icd, and the overexpressed AceK and AceA (Fig. 3), the decarboxylation of isocitrate by Icd could be bypassed through the glyoxylate shunt to reduce CO_2_ leakage. In our results, however, M+1 and M+3 AKG were increased in the adapted *E. coli* (Fig. 6). The increased M+1 AKG suggests that the decarboxylation of isocitrate by Icd was the main way to produce AKG, which is the essential precursor for glutamate and other derived amino acids, and thus cannot be bypassed completely. The increased M+3 AKG requires the availability of its precursor M+4 isocitrate, which required the participation of the ^13^C accumulated metabolites (Fig. 5), and thus implying the connection between the Asp-Thr cycle and the citrate cycle. When entering the glyoxylate shunt, isocitrate would be split into succinate and glyoxylate by the overexpressed AceA. However, the ^13^C enrichment changes of succinate had no statistical significancewere non-significantly increased in the adapted *E. coli*. Combining with the non-significant ^13^C enrichment changes of fumarate and malate (Fig. 6), which are the downstream metabolites of succinate, it is possible that there was an alternative pathway that drew succinate out of the citrate cycle. According to the dufferential expression of the adapted *E. coli* in autotrophic mode, SdhB and SucCD were downregulated (Fig. 3), which are responsible for converting succinate to fumarate and succinyl-CoA, respectively. In contrast, the overexpressed PrpBCD drove the reversed methylcitrate pathway (35), and thus bypassing the syntheses of fumarate and malate (Fig. 3). Therefore, the non-significant ^13^C enrichment results in succinate, fumarate, and malate indicate the competition of succinate between the citrate cycle and the reversed methylcitrate pathway (Fig. 6).

In summary, the increased M+1 pyruvate and M+1 lactate confirm the CO_2_ fixation by YdbK. The increased M+2 citrate derived from M+2 OAA confirms the CO_2_ fixation by Pck, and the main way to process OAA was citrate synthesis rather than malate conversion. The increased M+3 citrate implies the operation of the Asp-Thr cycle that provided ^13^C accumulated precursors. The increased M+1 AKG derived from M+2 OAA confirms the CO_2_ fixation by Pck, and the necessity of the decarboxylation by Icd to allow glutamate synthesis. The increased M+3 AKG implies the operation of the Asp-Thr cycle that provided ^13^C accumulated precursors. The non-significant results of succinate, fumarate, and malate indicate the operation of the methylcitrate pathway. Combining the results above, the integration of the carbon fixation module, the CGM cycle, and the Asp-Thr cycle into an autotrophic network is confirmed (Fig. 4).

## DISCUSSION

This study started with doubts about defining *E. coli* as a strict heterotroph, and the hypothesis that with enough time, *E. coli* could reach autotrophy with its native carbon fixation routes. It took 27 cumulative days of inorganic cultivation for the first doubling to emerge, manifesting its metabolic flexibility to develop autotrophy. Such basic feature could be overlooked due to insufficient adaptation time. After 1,000 days of consecutive inorganic subculturing, the autotrophic growth of the adapted *E. coli* reached 1,600 folds, and its full potential remains unknown, awaiting to be approached with more evolution time.

To understand how *E. coli* adapted its way to autotrophy, their accumulated mutations were examined, and *icd* (Asp398Glu) and *aceK* (Tyr474Ser) missense mutations are related to the metabolic switch between the citrate cycle and the glyoxylate shunt, which are most relevant to autotrophy. In previous study, three mutations were sufficient for their engineered *E. coli* to reach autotrophy, which occurred in genes phosphoglucoisomerase (*pgi*), RNA polymerase beta subunit (*rpoB*), and cAMP receptor protein (*crp*) (6), yet none of these genes was mutated in our results. There are five genes that were mutated in both previous and our studies, including *aceK*, *eutG*, succinyl-diaminopimelate desuccinylase (*dapE*), nitrate reductase Z subunit beta (*narY*), and lipopolysaccharide 3-alpha-galactosyltransferase (*waaO*), yet none of these shared mutated genes had the same mutation. For *aceK*, Tyr474Ser missense mutation was found in our results, whereas in previous studies, *aceK* was one of the 17 discarded genes in a large chromosomal deletion (4, 6). Such diverse mutation patterns show that there is no universal way for *E. coli* to reach autotrophy. Despite mutations that changed amino acid sequences and protein lengths were identified, more experiments are needed to clarify their effects.

Additionally, the roles of non-coding region mutations and synonymous mutations remain elusive (53, 54). Some non-coding region mutations might affect their target genes from more than 15kb away (55). Some synonymous mutations would affect gene expressions due to codon bias, or affect translation rates due to altered mRNA structure (56, 57). In terms of epigenetics, other than the mutations that changed methylation sites, the pattern of methylation could be more influential to phenotype. As a heritable yet flexible way of adaptation, epigenetic state is another form of selection (58), and the relations between methylome and autotrophy await to be investigated.

To identify the pathways taken by the adapted *E. coli* during inorganic cultivation, transcriptomic comparison was used to distinguish the key mechanisms for autotrophy. As a result, two CO_2_ fixing enzymes were found upregulated, namely pyruvate:ferredoxin oxidoreductase and PEP carboxykinase, which constitute the carbon fixation module as the shared root for the Asp-Thr cycle and the CGM cycle. These CO_2_ fixing enzymes originally serve as the replenishment mechanisms for the citrate cycle metabolites (59), and they are widely distributed among bacteria (60, 61). Such ubiquity suggests that some heterotrophs might have some capacity to fix CO_2_ (62, 63), and with enough time, they might develop autotrophic growth as well.

To track how CO_2_ was incorporated and converted in the adapted *E. coli* during autotrophic growth, ^13^CO_2_ labeling was applied. By comparing the ^13^C enrichment patterns of the citrate cycle metabolites, the favorable directions of the carbon fixation module and the CGM cycle were determined, the participations of the ^13^C accumulated metabolites from the Asp-Thr cycle were detected, and the integrated autotrophic network was confirmed. To further support this model, the ^13^C enrichment patterns of the methylcitrate cycle metabolites and amino acids can be analyzed.

After understanding the metabolic pathways taken by the adapted *E. coli* to reach autotrophy, the optimization of its carbon fixation efficiency can be designed rationally, by using mechanisms that are compatible with the autotrophic network. Firstly, to maximize the amount of fixed CO_2_, pyruvate:ferredoxin oxidoreductase, PEP carboxykinase, and other similar enzymes are reasonable targets (64–66). Another CO_2_ fixing enzyme worth considering is AKG oxidoreductase (Kor) (67, 68), which produces AKG and decreases the dependence on Icd that causes CO_2_ leakage. Secondly, to facilitate carbon fixation by accelerating the turnover rate of the autotrophic network, the downregulated enzymes in the Asp-Thr cycle can be overexpressed, and the overexpressed enzymes in the CGM cycle can be replaced with heterologous ones with higher efficiency. Thirdly, the autotrophic network can be reinforced by recycling its products, such as glyoxylate and propionyl-CoA. The recycling of glyoxylate can be strengthened by overexpressing GarR and Eno enzymes in the replenishment side loop of the Asp-Thr cycle (26), or overexpressing AceB and GlcB enzymes to enhance malate production, which could be converted to OAA as the shared precursor for the Asp-Thr cycle and the CGM cycle. The recycling of propionyl-CoA can be improved by introducing propionyl-CoA carboxylase (Pcc) that fixes bicarbonate to produce D-methylmalonyl-CoA, and methylmalonyl-CoA epimerase (Mce) to produce L-methylmalonyl-CoA, followed by overexpressing the native methylmalonyl-CoA mutase (ScpA) to produce succinyl-CoA (26), and thus reinforcing the CGM cycle. To recycle glyoxylate and propionyl-CoA simultaneously, 2-hydroxyglutarate synthase (Hgs) and 2-hydroxyglutarate dehydrogenase (Hgdh) can be introduced (26) to produce AKG without CO_2_ leakage, and thus reinforcing the CGM cycle. Finally, to test the bioproduction capacity of the autotrophic network, propionyl-CoA as its end-product and a versatile precursor can be revalorized. Once the designed pathway is introduced, the adapted *E. coli* can be a platform to convert propionyl-CoA to alcohols, fatty acids, bioplastics, and medical polyketides (69).

## MATERIALS AND METHODS

### Bacterial strain, plasmid, and transformation

The bacterial strain used in this study was *Escherichia coli* BW25113 parental strain from the KEIO collection (National BioResource Project, Japan), the plasmid used to provide antibiotics resistance was pGETS118 from lab collection (70). pGETS118 was introduced to *E. coli* BW25113 competent cells by heat shock transformation.

For competent cell chemical induction, 0.1 mL of *E. coli* BW25113 glycerol stock was inoculated in 5 mL of Luria-Bertani (LB) medium, cultivated in a roller drum at 120 rpm within an incubator (37°C), and the optical density at 600 nm (OD_600_) of bacterial culture was measured by the GeneQuant 1300 Spectrophotometer (GE Healthcare, USA) every 30 minutes. When OD_600_ reached 0.4, the bacterial culture was chilled on ice for 5 minutes. After centrifuging at 20,000 g (4°C) for 5 minutes, removed supernatant and resuspended bacterial pellet with 20 mL of 0.1 M MgCl_2_ (4°C). After centrifuging at 20,000 g (4°C) for 5 minutes, removed supernatant and resuspended bacterial pellet with 20 mL of 0.1 M CaCl_2_ (4°C), then chilled on ice for 20 minutes. After centrifuging at 20,000 g (4°C) for 5 minutes, removed supernatant and resuspended bacterial pellet with 20 mL of 0.1 M CaCl_2_ (4°C), chilled on ice for 10 minutes, and *E. coli* BW25113 competent cells were acquired.

For heat-shock transformation, 100 *µ*L of competent cells was gently mixed with 1 *µ*L of pGETS118 DNA solution (>100 ng), which was extracted from lab collection of *E. coli* with pGETS118 by the Plasmid Miniprep Purification Kit (GeneMark, USA), then chilled on ice for 20 minutes. Next, the mixed sample was heated in water bath (42°C) for 90 seconds, then chilled on ice for 20 minutes. To restore bacteria from heat damage, added LB medium 900 *µ*L to the mixed sample, and cultivated in a roller drum at 120 rpm within an incubator (37°C) for 2 hours. To select transformed bacteria, spread 0.1 mL of the restored sample on LB agar plate with 100 mg/L ampicillin, then placed up-side down in an incubator (37°C). After cultivating overnight, single colonies on plate were picked and inoculated in 5 mL of LB medium with 100 mg/L ampicillin. After cultivating overnight, LB cultures of the initial *E. coli* BW25113 with pGETS118 were acquired.

### Cultivation conditions

The base of M9 medium was composed of 47.8 mM Na_2_HPO_4_, 22 mM KH_2_PO_4_, 8.6 mM NaCl, and 18.7 mM NH_4_Cl. After autoclaving and cooling down, inorganic M9 medium was supplemented with solutions that were filter-sterilized through 0.22 *µ*m PVDF filters, reaching the final concentrations of 2 mM MgSO_4_, 100 *µ*M CaCl_2_, 100 mg/L ampicillin, and 36 *µ*M FeSO_4_ from freshly prepared filter-sterilized FeSO_4_ solution to avoid Fe^2+^ oxidation overtime. For organic M9 medium, a filter-sterilized glucose solution was supplemented additionally, reaching a final concentration of 0.125 % (w/v) glucose. For small volume cultivation of 5 mL, Hungate tubes were used, leaving 10 mL of headspace volume. For larger volume cultivation of 150 mL, crimp-top bottles were used, leaving 100 mL of headspace volume. Containers were closed with rubber stoppers after medium distribution and bacteria inoculation, and the partial pressure of CO_2_ was elevated to 10% by injecting CO_2_ through sterilized syringe. For cultivation, Hungate tubes were placed in a roller drum at 120 rpm within an incubator (37°C), and crimp-top bottles were placed in a shaking incubator (37°C) at 200 rpm.

### Growth curve measurement and ALE experiments

For growth curve measurement, 0.2 mL of inorganic culture from each tube was extracted daily by sterilized syringe, and 0.1 mL of the extracted culture was serially diluted, spread evenly on LB agar plate with 100 mg/L ampicillin by autoclaved glass beads, then placed up-side down in an incubator (37°C). After cultivating overnight, viable bacterial numbers were determined by counting CFU. To calculate CFU/mL, the values of CFU were first multiplied by ten to scale up the CFU from 0.1 mL to 1 mL, then multiplied by their dilution factors. To calculate growth fold-change, the CFU/mL at hour N was divided by the inoculated CFU/mL.

To prepare for ALE, the LB culture of the initial *E. coli* BW25113 was washed three times by repeating the cycle of centrifugation, supernatant removal, and pellet resuspension with inorganic M9 medium, in order to remove residual LB medium that would interfere inorganic cultivation. Then, the washed culture was serially diluted to 1:10,000 with inorganic M9 medium, and six diluted inoculums were acquired.

For the first phase of ALE, 0.5 mL of each diluted inoculum was inoculated in 4.5 mL of inorganic M9 medium. After inorganic cultivation for 72 hours, 1 mL of each remaining culture was transferred to 4 mL of organic M9 medium with 0.125 % (w/v) glucose, reaching a final concentration of 0.1 % (w/v) glucose. After organic cultivation for 24 hours, the repopulated cultures were washed three times and serially diluted to 1:1,000, and 0.5 mL of each diluted inoculum was inoculated in 4.5 mL of inorganic M9 medium, then cultivated for 72 hours. By repeating the steps above, the inorganic-organic rotation continued.

The second phase of ALE began when the growth fold-change of inorganic cultivation in the first phase reached 10 folds. Inorganic cultures of the adapted *E. coli* were washed three times, diluted proportionally, and 0.5 mL of each inoculum was subcultured in 4.5 mL of inorganic M9 medium. Before growth fold-change reached 1,000 folds, the subculturing time point was at 96 hours, and the remaining culture was kept cultivating until 192 hours. After growth fold-change reached 1,000 folds, the subculturing time point was changed to 192 hours. By repeating the steps above, the consecutive inorganic subculturing continued. Proportional dilution rate for subculturing was adjusted by the growth fold-change of each batch. When reaching growth fold-changes of 100 and 1,000 folds, dilution rates for subculturing were adjusted to 1:10 and 1:100, then another 1:10 dilution was done by subculturing process. To avoid growth fold-change overestimation or underestimation that would be caused by lower or higher inoculated CFU/mL, dilution rates for subculturing were further calibrated by the CFU/mL values 24 hours prior to subculturing, and the batches with 1,500–2,500 inoculated CFU/mL were analyzed.

### DNA sample collection, sequencing, and mutation analysis

For sample cultivation, the adapted *E. coli* after 1,000 days of consecutive inorganic subculturing was subcultured in inorganic M9 medium. To increase the chance of picking the clones that could grow autotrophically, the inorganic culture had to be sampled before 72 hours, which was the first peaking time of its population. After inorganic cultivation for 48 hours, 0.1 mL of the inorganic culture was spread on LB agar plate with 100 mg/L ampicillin. After cultivating overnight, five single colonies were randomly picked and inoculated in LB medium with 100 mg/L ampicillin. After cultivating overnight, five cultures of the adapted *E. coli* clones were acquired.

For DNA extraction, the Wizard Genomic DNA Purification Kit (Promega, USA) was used. To assess DNA purity, quantity, and integrity, the NanoDrop 2000c Spectrophotometer (Thermo Fisher Scientific, USA), the Qubit 4.0 fluorometer (Thermo Fisher Scientific, USA) with the Quant-iT dsDNA High-Sensitivity Assay Kit (Thermo Fisher Scientific, USA), and the Fragment Analyzer (Agilent Technologies, USA) were used. For DNA fragmentation, 200 ng of DNA from each clone was fragmented to a major peak size range of 350–400 bp by the M220 Focused-ultrasonicator (Covaris, USA). For library construction, the SureSelect XT Low Input Reagent Kit (Agilent, USA) was used, and the fragmented DNA was ligated with the Illumina TruSeq Unique Dual Indexes (Integrated DNA Technologies, USA), then amplified by PCR for 10 cycles. For library purification, the AMPure XP beads (Beckman Coulter, USA) were used. For size selection, the purified libraries were selected for 350–600 bp fragments by the BluePippin DNA Size Selection System (Sage Science, USA). To assess library purity, quantity, and size distribution, the NanoDrop 2000c Spectrophotometer (Thermo Fisher Scientific, USA), the Qubit 4.0 fluorometer (Thermo Fisher Scientific, USA) with the Quant-iT dsDNA High-Sensitivity Assay Kit (Thermo Fisher Scientific, USA), and the 4200 TapeStation System (Agilent, USA) with the D1000 ScreenTape Assay (Agilent, USA) were used. DNA sequencing was performed on the Illumina NovaSeq X Series (Illumina, USA), using the NovaSeq X Series 25B Reagent 300 cycle Kit (Illumina, USA) by the 150 bp paired-end protocol.

For read qualification, FASTQ reads were processed through Trimmomatic v0.36 (71) for adapter removal, quality trimming, and length filtering. For sequence alignment, qualified reads were aligned with *E. coli* BW25113 genome reference (17) and pGETS118 plasmid reference (72) by BWA-MEM v0.7.17 (73). For variant identification, Genome Analysis Toolkit v4.1.9 (74) was applied. For variant annotation, SnpEff v4.3t (75) was applied.

### RNA sample collection, sequencing, and transcriptome analysis

For sample cultivation in larger volume, the adapted *E. coli* was cultivated in crimp-top bottle with 150 mL of medium. For autotrophic mode bacterial samples, to acquire sufficient viable bacterial number for transcriptome analysis, 18 bottles with inorganic M9 medium were used to cultivate the adapted *E. coli*, and were collected after 48 hours of cultivation to capture their transcription profiles in their growing phase. For heterotrophic mode bacterial samples, two bottles with organic M9 medium were used to cultivate the adapted *E. coli*, and were collected after 2 hours to capture their transcription profiles in their growing phase, and also to match the CFU/mL level of autotrophic mode samples, avoiding transcriptional changes caused by different densities of bacteria. For sample collection, the adapted *E. coli* cultures of different modes were vacuum filtered separately through sterilized 0.22 *µ*m PVDF membrane, and bacteria on filter was washed down by 1 mL of corresponding M9 medium (4°C). After centrifuging at 20,000 g for 10 minutes (4°C) and removing supernatant, two tubes of autotrophic mode bacterial samples and two tubes of heterotrophic mode bacterial samples were acquired.

For RNA extraction, the Invitrogen TRIzol Plus RNA Purification Kit (Thermo Fisher Scientific, USA) was used. To assess RNA purity and quantity, the NanoDrop 1000 Spectrophotometer (Thermo Fisher Scientific, USA), and the Bioanalyzer 2100 (Agilent Technology, USA) with the RNA 6000 Nano Kit (Agilent Technology, USA) were used. For rRNA depletion, 0.5 µg of RNA from each tube was processed through the Invitrogen RiboMinus Bacteria 2.0 Transcriptome Isolation Kit (Thermo Fisher Scientific, USA), quantified by NanoDrop 1000 Spectrophotometer (Thermo Fisher Scientific, USA), then assessed size distribution by the Bioanalyzer 2100 (Agilent Technology, USA) with the RNA 6000 Nano Kit (Agilent Technology, USA).

For RNA library construction, the SureSelect XT HS2 mRNA Library Preparation Kit (Agilent, USA) was used, and 10 uL of the rRNA depleted sample from each tube was ligated with the Unique Dual Indexes (Agilent, USA), then amplified by PCR for 10 cycles. For library purification, the HighPrep PCR Clean-Up System (MagBio Genomics, USA) was used. To assess library purity, quantity, and size distribution, the NanoDrop 2000c Spectrophotometer (Thermo Fisher Scientific, USA), the Qubit 4.0 fluorometer (Thermo Fisher Scientific, USA) with the Quant-iT dsDNA High-Sensitivity Assay Kit (Thermo Fisher Scientific, USA), and the 4200 TapeStation System (Agilent, USA) with the D1000 ScreenTape Assay (Agilent, USA) were used. RNA sequencing was performed on the Illumina NovaSeq X Series (Illumina, USA), using the NovaSeq X Series 25B 300 cycle Reagent Kit (Illumina, USA) by the 150 bp paired-end protocol.

To generate FASTQ reads as sequencing data, BCL Convert v4.2.4 (Illumina, USA) was applied. For read qualification, FASTQ reads were processed through Trimmomatic v0.36 (71) for adapter removal, quality trimming, and length filtering. For sequence alignment, qualified reads were aligned with genome reference by HISAT2 (76). For transcript assembly and quantification, StringTie was used (76), and gene expression levels were normalized to transcripts per million (TPM) (77). For differential expression (DE) analysis, StringTie v2.1.7, DESeq v1.38.0, and DESeq2 v1.38.0 (78) with genome bias detection and correction were applied. To calculate expression fold-change, the TPM of the adapted *E. coli* in autotrophic mode was divided by the TPM of the adapted *E. coli* in heterotrophic mode. To assess statistical significance, p-values were derived from composite null hypothesis testing, and q-values were the multiple testing adjusted p-values calculated by the Benjamini-Hochberg method (79).

### ^13^CO_2_ labeling, and ^13^C enrichment analysis of the citrate cycle metabolites

For larger volume of cultivation, bacterial strains were cultivated in the crimp-top bottles with 150 mL of inorganic M9 medium. For ^13^CO_2_ labeling, the partial pressure of ^13^CO_2_ was elevated to 10% by injecting ^13^CO_2_ through sterilized syringe. To have sufficient viable bacteria for ^13^C enrichment analysis of the citrate cycle metabolites, two bottles of the adapted *E. coli* and 18 bottles of the initial *E. coli* were cultivated. For bacterial sample collection, inorganically cultivated *E. coli* of different adaptation phases were vacuum filtered separately through sterilized 0.22 *µ*m PVDF membrane, and bacteria on filter was washed down by 1 mL of inorganic M9 medium (4°C). After centrifuging at 20,000 g for 10 minutes (4°C) and removing supernatant, two tubes of the adapted *E. coli* pellets and two tubes of the initial *E. coli* pellets were acquired, then stored in a freezer (−80°C) until ^13^C enrichment analysis.

For sample preparation and analysis, the protocol from previous study was applied (68). For sample quenching to arrest metabolism, 400 *µ*L methanol (−20°C) and 200 *µ*L distilled water (4°C) were added to each pelleted sample, then homogenized with glass rod and sonicated in ultrasonic bath (4°C) for 5 minutes. For sample extraction, added 400 *µ*L chloroform to each quenched sample, vortexed thoroughly, then shook on a shaker at 200 rpm in a fridge (4°C) for 10 minutes. After centrifuging at 14,000 g (4°C) for 10 minutes, the upper phase containing polar metabolites from each sample was transferred to a new tube, then vacuum dried (55°C) for 2 hours. For methoximation of pyruvate and AKG, each dried sample was resuspended in 30 *µ*L pyridine with 10 mg/mL methoxyaime hydrochloride (4°C), vortexed thoroughly and span down, then heated in dry bath (73°C) for 30 minutes. For derivatization, 20 *µ*L N-tert-butyldimethylsilyl-N-methyltrifluoroacetamide (MTBSTFA, 4°C) was added to each methoximated sample, vortexed thoroughly and span down, then heated in dry bath (73°C) for 1 hour. After centrifuging at 14,000 g (4°C) for 5 minutes, derivatized samples were acquired. For GC-MS analysis, each derivatized sample was separated in a DB-225MS 30 m × 0.25 mm column (Agilent Technologies, USA), and analyzed in electron ionization mode by Model 6890 Gas Chromatograph (Agilent Technologies, USA) and Model 5973 Mass Spectrometer (Agilent Technologies, USA).

The comparison of fractional ^13^C enrichment mean values between the adapted and the initial *E. coli* were analyzed by t-test, and statistical significance was defined as value < 0.05, borderline significance was defined as p-value < 0.1.

## ACKNOWLEDGMENTS

This work was funded by National Science and Technology Council (NSTC 113-2221-E-005-020-MY3). We thank Wei-Lun Guo for technical support in the early stage. We thank Welgene Biotech Co., Ltd. (Taiwan) for sequencing services.

Conceptualization: Shao-Yu Huang and Chieh-Chan Huang; methodology: Shao-Yu Huang, Jian-Hau Peng, and Yu-Hsi Lin; investigation: Shao-Yu Huang, Jian-Hau Peng, Shou-Chen Lo, and Chia-Ho Liu; visualization: Shao-Yu Huang; writing—original manuscript: Shao-Yu Huang; writing—review and editing: Shao-Yu Huang, Jian-Hau Peng, Yu-His Lin, En-Pei Isabel Chiang, and Chieh-Chen Huang; supervision: En-Pei Isabel Chiang and Chieh-Chen Huang. All authors reviewed and approved the final manuscript.

## REFERENCES

1. Forster PM, Smith C, Walsh T, Lamb WF, Lamboll R, Cassou C, Hauser M, Hausfather Z, Lee J-Y, Palmer MD, Von Schuckmann K, Slangen ABA, Szopa S, Trewin B, Yun J, Gillett NP, Jenkins S, Matthews HD, Raghavan K, Ribes A, Rogelj J, Rosen D, Zhang X, Allen M, Aleluia Reis L, Andrew RM, Betts RA, Borger A, Broersma JA, Burgess SN, Cheng L, Friedlingstein P, Domingues CM, Gambarini M, Gasser T, Gütschow J, Ishii M, Kadow C, Kennedy J, Killick RE, Krummel PB, Liné A, Monselesan DP, Morice C, Mühle J, Naik V, Peters GP, Pirani A, Pongratz J, Minx JC, Rigby, Matthew, Rohde R, Savita A, Seneviratne SI, Thorne P, Wells C, Western LM, van der Werf GR, Wijffels SE, Masson-Delmotte V, Zhai PM. 2025. Indicators of Global Climate Change 2024: Annual update of key indicators of the state of the climate system and human influence. Earth Syst Sci Data 17:2641–2680.

2. Claassens NJ, Sousa DZ, dos Santos VAPM, de Vos WM, van der Oost J. 2016. Harnessing the power of microbial autotrophy. Nat Rev Microbiol 14:692–706.

3. Pontrelli S, Chiu TY, Lan EI, Chen FYH, Chang P, Liao JC. 2018. *Escherichia coli* as a host for metabolic engineering. Metab Eng 50:16–46.

4. Gleizer S, Ben-Nissan R, Bar-On YM, Antonovsky N, Noor E, Zohar Y, Jona G, Krieger E, Shamshoum M, Bar-Even A, Milo R. 2019. Conversion of *Escherichia coli* to generate all biomass carbon from CO_2_. Cell 179:1255–1263.

5. Antonovsky N, Gleizer S, Noor E, Zohar Y, Herz E, Barenholz U, Zelcbuch L, Amram S, Wides A, Tepper N, Davidi D, Bar-On Y, Bareia T, Wernick David G, Shani I, Malitsky S, Jona G, Bar-Even A, Milo R. 2016. Sugar synthesis from CO_2_ in *Escherichia coli*. Cell 166:115–125.

6. Ben-Nissan R, Milshtein E, Pahl V, de Pins B, Jona G, Levi D, Yung H, Nir N, Ezra D, Gleizer S, Link H, Noor E, Milo R. 2024. Autotrophic growth of *E. coli* is achieved by a small number of genetic changes. eLife 12:RP88793.

7. Satanowski A, Dronsella B, Noor E, Vögeli B, He H, Wichmann P, Erb TJ, Lindner SN, Bar-Even A. 2020. Awakening a latent carbon fixation cycle in *Escherichia coli*. Nat Commun 11:5812.

8. Blount ZD, Borland CZ, Lenski RE. 2008. Historical contingency and the evolution of a key innovation in an experimental population of *Escherichia coli*. Proc Natl Acad Sci USA 105:7899–7906.

9. Lenski RE. 2017. Experimental evolution and the dynamics of adaptation and genome evolution in microbial populations. ISME J 11:2181–2194.

10. Malpica R, Franco B, Rodriguez C, Kwon O, Georgellis D. 2004. Identification of a quinone-sensitive redox switch in the ArcB sensor kinase. Proc Natl Acad Sci USA 101:13318–13323.

11. Doyle SA, Beernink PT, Koshland DE. 2001. Structural basis for a change in substrate specificity: Crystal structure of S113E isocitrate dehydrogenase in a complex with isopropylmalate, Mg^2+^, and NADP. Biochem 40:4234–4241.

12. Li QJ, Zheng JM, Tan HW, Li XC, Chen GJ, Jia ZC. 2013. Unique kinase catalytic mechanism of AceK with a single magnesium ion. PLoS One 8:e72048.

13. Zhang XY, Shen QY, Lei Z, Wang QY, Zheng JM, Jia ZC. 2019. Characterization of metal binding of bifunctional kinase/phosphatase AceK and implication in activity modulation. Sci Rep 9:9198.

14. Fischer E, Sauer U. 2003. A novel metabolic cycle catalyzes glucose oxidation and anaplerosis in hungry *Escherichia coli*. J Biol Chem 278:46446–46451.

15. Sánchez-Romero MA, Casadesús J. 2019. The bacterial epigenome. Nat Rev Microbiol 18:7–20.

16. Westphal LL, Sauvey P, Champion MM, Ehrenreich IM, Finkel SE. 2016. Genomewide Dam methylation in *Escherichia coli* during long-term stationary phase. mSystems 1:e00130–16.

17. Grenier F, Matteau D, Baby V, Rodrigue S. 2014. Complete genome sequence of *Escherichia coli* BW25113. Genome Announc 2:e01038–14.

18. Ren J, Wang W, Nie JL, Yuan WQ, Zeng AP. 2022. Understanding and engineering glycine cleavage system and related metabolic pathways for C1-based biosynthesis. Adv Biochem Eng Biotechnol 180:273–298.

19. Yishai O, Goldbach L, Tenenboim H, Lindner SN, Bar-Even A. 2017. Engineered assimilation of exogenous and endogenous formate in *Escherichia coli*. ACS Synth Biol 6:1722–1731.

20. Bang J, Hwang CH, Ahn JH, Lee JA, Lee SY. 2020. *Escherichia coli* is engineered to grow on CO_2_ and formic acid. Nat Microbiol 5:1459–1463.

21. Aslan S, Noor E, Bar-Even A. 2017. Holistic bioengineering: Rewiring central metabolism for enhanced bioproduction. Biochem J 474:3935–3950.

22. Zhang XL, Jantama K, Moore JC, Jarboe LR, Shanmugam KT, Ingram LO. 2009. Metabolic evolution of energy-conserving pathways for succinate production in *Escherichia coli*. Proc Natl Acad Sci USA 106:20180–20185.

23. Nakayama T, Shin-Ichiro Y, Yonei S, Zhang-Akiyama Q-M. 2013. *Escherichia coli* pyruvate:flavodoxin oxidoreductase, YdbK - regulation of expression and biological roles in protection against oxidative stress. Genes Genet Syst 88: 175–188.

24. Akhtar MK, Jones PR. 2009. Construction of a synthetic YdbK-dependent pyruvate:H2 pathway in *Escherichia coli* BL21(DE3). Metab Eng 11:139–147.

25. Schulz-Mirbach H, Müller A, Wu T, Pfister P, Aslan S, von Borzyskowski LS, Erb TJ, Bar-Even A, Lindner SN. 2022. On the flexibility of the cellular amination network in *E coli*. eLife 11:e77492.

26. Bar-Even A, Noor E, Lewis NE, Milo R. 2010. Design and analysis of synthetic carbon fixation pathways. Proc Natl Acad Sci USA 107:8889–8894.

27. Muchowska KB, Varma SJ, Moran J. 2020. Nonenzymatic metabolic reactions and life’s origins. Chem Rev 120:7708–7744.

28. Gerike U, Hough DW, Russell NJ, Dyall-Smith ML, Danson MJ. 1998. Citrate synthase and 2-methylcitrate synthase: structural, functional and evolutionary relationships. Microbiology 144:929–935.

29. Blank L, Green J, Guest JR. 2002. AcnC of *Escherichia coli* is a 2-methylcitrate dehydratase (PrpD) that can use citrate and isocitrate as substrates. Microbiology 148:133–146.

30. Yang P, Liu WJ, Chen YA, Gong AD. 2022. Engineering the glyoxylate cycle for chemical bioproduction. Front Bioeng Biotechnol 10:1066651.

31. Peng J, Sun JR, Luo YC, Wu H. 2025. Evolution-assisted engineering of *E. coli* improved succinic acid production from glycerol. BioDesign Res 7:100022.

32. Cronan JE, Laporte D. 2005. Tricarboxylic acid cycle and glyoxylate bypass. EcoSal Plus 1:10.1128/ecosalplus.3.5.2.

33. Horswill AR, Escalante-Semerena JC. 1999. The *prpE* gene of *Salmonella typhimurium* LT2 encodes propionyl-CoA synthetase. Microbiology 145:1381–1388.

34. van der Rest ME, Christian F, Douwe M. 2000. Functions of the membrane-associated and cytoplasmic malate dehydrogenases in the citric acid cycle of *Escherichia coli*. J Bacteriol 182:6892–6899.

35. Serafini A, Tan L, Horswell S, Howell S, Greenwood DJ, Hunt DM, Phan MD, Schembri M, Monteleone M, Montague CR, Britton W, Garza-Garcia A, Snijders AP, VanderVen B, Gutierrez MG, West NP, de Carvalho LPS. 2019. *Mycobacterium tuberculosis* requires glyoxylate shunt and reverse methylcitrate cycle for lactate and pyruvate metabolism. Mol Microbiol 112:1284–1307.

36. Cotton CA, Bernhardsgrütter I, He H, Burgener S, Schulz L, Paczia N, Dronsella B, Erban A, Toman S, Dempfle M, De Maria A, Kopka J, Lindner SN, Erb TJ, Bar-Even A. 2020. Underground isoleucine biosynthesis pathways in *E. coli*. eLife 9:e54207.

37. Janßen HJ, Steinbüchel A. 2014. Fatty acid synthesis in *Escherichia coli* and its applications towards the production of fatty acid based biofuels. Biotechnol Biofuels 7:7.

38. Bak DW, Weerapana E. 2023. Monitoring Fe–S cluster occupancy across the *E. coli* proteome using chemoproteomics. Nat Chem Biol 19:356–366.

39. Mey AR, Gómez-Garzón C, Payne SM. 2021. Iron transport and metabolism in *Escherichia*, *Shigella*, and *Salmonella*. EcoSal Plus 9:eESP-0034-2020.

40. Tsylents U, Burmistrz M, Wojciechowska M, Stępień J, Maj P, Trylska J. 2024. Iron uptake pathway of *Escherichia coli* as an entry route for peptide nucleic acids conjugated with a siderophore mimic. Front Microbiol 15:1331021.

41. Grinter R, Lithgow T. 2019. The structure of the bacterial iron–catecholate transporter Fiu suggests that it imports substrates via a two-step mechanism. J Biol Chem 294:19523–19534.

42. van der Ploeg JR, Eichhorn E, Leisinger T. 2001. Sulfonate-sulfur metabolism and its regulation in *Escherichia coli*. Arch Microbiol 176:1–8.

43. Esquilin-Lebron K, Dubrac S, Barras F, Boyd JM. 2021. Bacterial approaches for assembling iron-sulfur proteins. mBio 12:e02425–21.

44. Ren XJ, Liang F, He ZF, Fan BQ, Zhang ZR, Guo XD, Du YK, Pang YL, Li JH, Lyu JX, Tan GQ. 2021. Identification of an intermediate form of ferredoxin that binds only iron suggests that conversion to holo-ferredoxin is independent of the ISC system in *Escherichia coli*. Appl Environ Microbiol 87:e03153–20.

45. Corless EI, Mettert EL, Kiley PJ, Antony E. 2019. Elevated expression of a functional Suf pathway in *Escherichia coli* BL21(DE3) enhances recombinant production of an iron-sulfur cluster-containing protein. J Bacteriol 202:10.1128/jb.00496-19.

46. Blahut M, Sanchez E, Fisher CE, Outten FW. 2020. Fe-S cluster biogenesis by the bacterial Suf pathway. Biochim Biophys Acta Mol Cell Res 1867:118829.

47. Schwartz CJ, Giel JL, Patschkowski T, Luther C, Ruzicka FJ, Beinert H, Kiley PJ. 2001. IscR, an Fe-S cluster-containing transcription factor, represses expression of *Escherichia coli* genes encoding Fe-S cluster assembly proteins. Proc Natl Acad Sci USA 98:14895–14900.

48. Dai YY, Outten FW. 2012. The *E. coli* SufS - SufE sulfur transfer system is more resistant to oxidative stress than IscS - IscU. FEBS lett 586:4016–4022.

49. Boyd ES, Thomas KM, Dai YY, Boyd JM, Outten FW. 2014. Interplay between oxygen and Fe–S cluster biogenesis: Insights from the Suf pathway. Biochemistry 53:5834–5847.

50. Liu YLA, Lee CC, Górecki K, Stiebritz MT, Duffin C, Solomon JB, Ribbe MW, Hu YL. 2025. Heterologous synthesis of a simplified nitrogenase analog in *Escherichia coli*. Sci Adv 11:eadw6785.

51. Wang QZ, Ou MS, Kim Y, Ingram LO, Shanmugam KT. 2010. Metabolic flux control at the pyruvate node in an anaerobic *Escherichia coli* strain with an active pyruvate dehydrogenase. Appl Environ Microbiol 76:2107–2114.

52. Lindeque JZ. 2024. Targeted analysis of organic acids with GC-MS/MS: Challenges and prospects. Anal Biochem 694:115620.

53. Lamoureux CR, Phaneuf PV, Palsson Bernhard O, Zielinski Daniel C. 2024. Escherichia coli non-coding regulatory regions are highly conserved. NAR Genom Bioinform 6:lqae041.

54. Bailey SF, Hinz A, Kassen R. 2014. Adaptive synonymous mutations in an experimentally evolved Pseudomonas fluorescens population. Nat Commun 5:4076.

55. Bondarenko V, Liu Y, Ninfa A, Studitsky VM. 2002. Action of prokaryotic enhancer over a distance does not require continued presence of promoter-bound *σ*54 subunit. Nucleic Acids Res 30:636–642.

56. Yang DD, Rusch LM, Widney KA, Morgenthaler AB, Copley SD. 2024. Synonymous edits in the *Escherichia coli* genome have substantial and condition-dependent effects on fitness. Proc Natl Acad Sci USA 121:e2316834121.

57. Bailey SF, Alonso Morales LA, Kassen R. 2021. Effects of synonymous mutations beyond codon bias: The evidence for adaptive synonymous substitutions from microbial evolution experiments. Genome Biol Evol 13:evab141.

58. Beaulaurier J, Schadt EE, Fang G. 2018. Deciphering bacterial epigenomes using modern sequencing technologies. Nat Rev Genet 20:157–172.

59. Erb TJ. 2011. Carboxylases in natural and synthetic microbial pathways. Appl Environ Microbiol 77:8466–8477.

60. Duwor S, Brites D, Mäser P. 2024. Phylogenetic analysis of pyruvate-ferredoxin oxidoreductase, a redox enzyme involved in the pharmacological activation of nitro-based prodrugs in bacteria and protozoa. Biology 13:178.

61. Walker RP, Chen ZH, Famiani F. 2021. Gluconeogenesis in plants: A key interface between organic acid/amino acid/lipid and sugar metabolism. Molecules 26:5129.

62. Spona-Friedl M, Braun A, Huber C, Eisenreich W, Griebler C, Kappler A, Elsner M. 2020. Substrate-dependent CO_2_ fixation in heterotrophic bacteria revealed by stable isotope labelling. FEMS Microbiol Ecol 96:fiaa080.

63. Taubert M, Overholt WA, Heinze BM, Matanfack GA, Houhou R, Jehmlich N, von Bergen M, Rösch P, Popp J, Küsel K. 2021. Bolstering fitness via CO_2_ fixation and organic carbon uptake: Mixotrophs in modern groundwater. ISME J 16:1153–1162.

64. Bonitatibus SC, Pham RV, Weitz AC, Lopéz-Muñoz MM, Li B, Metcalf WW, Nair SK, Elliott SJ. 2025. The redox landscape of pyruvate:ferredoxin oxidoreductases reveals often conserved Fe–S cluster potentials. J Biol Chem 301:110380.

65. Zhu LW, Zhang L, Wei LN, Li HM, Yuan ZP, Chen T, Tang YL, Liang XH, Tang YJ. 2015. Collaborative regulation of CO_2_ transport and fixation during succinate production in *Escherichia coli*. Sci Rep 5:17321.

66. Park S, Lee JU, Cho S, Kim H, Oh HB, Pack SP, Lee J. 2017. Increased incorporation of gaseous CO_2_ into succinate by *Escherichia coli* overexpressing carbonic anhydrase and phosphoenolpyruvate carboxylase genes. J Biotechnol 241:101–107.

67. Lo SC, Chiang EPI, Yang YT, Li SY, Peng JH, Tsai SY, Wu DY, Yu CH, Huang CH, Su TT, Tsuge K, Huang CC. 2021. Growth enhancement facilitated by gaseous CO_2_ through heterologous expression of reductive tricarboxylic acid cycle genes in *Escherichia coli*. Fermentation 7:98.

68. Peng JH, Lo SC, Yu YN, Yang YT, Chen YC, Tsai AI, Wu DY, Huang CH, Su TT, Huang CC, Chiang EPI. 2025. Carbon fluxes rewiring in engineered *E. coli* via reverse tricarboxylic acid cycle pathway under chemolithotrophic condition. J Biol Eng 19:20.

69. Srirangan K, Bruder M, Akawi L, Miscevic D, Kilpatrick S, Moo-Young M, Chou CP. 2017. Recent advances in engineering propionyl-CoA metabolism for microbial production of value-added chemicals and biofuels. Crit Rev Biotechnol 37:701–722.

70. Chin WC, Lin KH, Liu CC, Tsuge K, Huang CC. 2017. Improved n-butanol production via co-expression of membrane-targeted tilapia metallothionein and the clostridial metabolic pathway in *Escherichia coli*. BMC Biotechnol 17:36.

71. Bolger AM, Lohse M, Usadel B. 2014. Trimmomatic: A flexible trimmer for Illumina sequence data. Bioinformatics 30:2114–2120.

72. Tsuge K, Sato Y, Kobayashi Y, Gondo M, Hasebe M, Togashi T, Tomita M, Itaya M. 2015. Method of preparing an equimolar DNA mixture for one-step DNA assembly of over 50 fragments. Sci Rep 5:10655.

73. Li H. 2013. Aligning sequence reads, clone sequences and assembly contigs with BWA-MEM. arXiv 10.48550/arXiv.1303.3997.

74. McKenna A, Hanna M, Banks E, Sivachenko A, Cibulskis K, Kernytsky A, Garimella K, Altshuler D, Gabriel S, Daly M, DePristo MA. 2010. The Genome Analysis Toolkit: A MapReduce framework for analyzing next-generation DNA sequencing data. Genome Res 20:1297–1303.

75. Cingolani P, Platts A, Wang LL, Coon M, Nguyen T, Wang LL, Land SJ, Lu Xy, Ruden DM. 2012. A program for annotating and predicting the effects of single nucleotide polymorphisms, SnpEff. Fly 6:80–92.

76. Pertea M, Kim D, Pertea GM, Leek JT, Salzberg SL. 2016. Transcript-level expression analysis of RNA-seq experiments with HISAT, StringTie and Ballgown. Nat Protoc 11:1650–1667.

77. Wagner GP, Kin K, Lynch VJ. 2012. Measurement of mRNA abundance using RNA-seq data: RPKM measure is inconsistent among samples. Theory Biosci 131:281–285.

78. Sahraeian SME, Mohiyuddin M, Sebra R, Tilgner H, Afshar PT, Au KF, Bani Asadi N, Gerstein MB, Wong WH, Snyder MP, Schadt E, Lam HYK. 2017. Gaining comprehensive biological insight into the transcriptome by performing a broad-spectrum RNA-seq analysis. Nat Commun 8:59.

79. Hochberg Y, Benjamini Y. 1990. More powerful procedures for multiple significance testing. Stat Med 9:811–818.

